# Left-Lateralized Contributions of Saccades to Cortical Activity during a One-Back Word Recognition Task

**DOI:** 10.1101/211177

**Authors:** Yu-Cherng C. Chang, Sheraz Khan, Samu Taulu, Gina Kuperberg, Emery N. Brown, Matti S. Hämäläinen, Simona Temereanca

## Abstract

Saccadic eye movements are an inherent component of natural reading, yet their contribution to information processing at subsequent fixation remains elusive. Here we use anatomically-constrained magnetoencephalography (MEG) to examine cortical activity following saccades as healthy human subjects engaged in a one-back word recognition task. This activity was compared with activity following external visual stimulation that mimicked saccades. A combination of procedures were employed to eliminate saccadic ocular artifacts from the MEG signal. Both saccades and saccade-like external visual stimulation produced early-latency responses beginning ~70 ms after onset in occipital cortex and spreading through the ventral and dorsal visual streams to temporal, parietal and frontal cortices. Robust differential activity following the onset of saccades versus similar external visual stimulation emerged during 150-350 ms in a left-lateralized cortical network. This network included (i) left lateral occipitotemporal and nearby inferotemporal cortex, (ii) left posterior Sylvian fissure and nearby multimodal cortex, and (iii) medial parietooccipital, posterior cingulate and retrosplenial cortices. Moreover, this left-lateralized network colocalized with word repetition priming effects. Together, results suggest that central saccadic mechanisms influence a left-lateralized language network in occipitotemporal and temporal cortex above and beyond saccadic influences at preceding stages of information processing during visual word recognition.

## 1. INTRODUCTION

Active reading is a complex skill thought to require coordination between eye movements, attention, and written language processing (Engbert et al., 2005; Pollatsek et al., 2006; Rayner, 2009), yet the neural basis of this coupling remains elusive. In particular, the extent to which saccades impact reading processes at fixation is little understood, since current neurobiological models of visual word processing typically account for neuroimaging and neurophysiological data collected during stable fixation when the natural temporal proximity of saccades and target words is disrupted (Sereno and Rayner, 2003). Recent studies using fMRI and EEG combined with eye tracking have begun to evaluate the ecological validity of these models in reading tasks that include eye movements, albeit focusing on mechanisms of eye movement control and contributions of spatial attention rather than the impact of the saccade itself on information processing at fixation (e.g., Dimigen et al., 2012; Henderson et al., 2015, 2016; Schuster et al., 2015, 2016; but see Kornrumpf et al., 2016). However, active vision studies in humans and monkey indicate that central saccadic signals gated by brain regions that control eye movements and attention modify visual perception and cognition around the time of saccades, acting via distinct mechanisms before, during and after an eye movement (Wurtz, 2008; Berman and Colby, 2009; Ibbotson and Krekelberg, 2011). Recent evidence using magnetoencephalography (MEG) combined with saccade detection in real time supports the view that saccades also impact brain responses to words presented at fixation, revealing various degrees of saccadic modulation in early visual and higher cortical areas (Temereanca et al., 2012). An unexplored question is whether central saccadic signals impact occipitotemporal and temporal cortical areas that are implicated in visual word processing above and beyond preceding saccadic influences in occipital cortex.

During saccades, visual stability emerges in the brain despite abrupt self-induced changes in visual input associated with the eye movement. In contrast to salient external visual motion, self-induced saccadic visual motion is not consciously perceived, although it continues to be processed in the visual system and thus can impact information processing at subsequent fixation (Ibbotson and Cloherty, 2009). Consistent with interactions between visual signals during and after saccades, our previous results suggest that saccadic image motion modulates responses to words presented at fixation (Temereanca et al., 2012). In addition to such visual effects, central saccadic signals mediated by brain regions that control eye movements and attention are known to impact information processing in visual areas, producing effects that include transsaccadic suppression followed by postsaccadic enhancement. Transaccadic suppression from ~100 ms before onset to ~50 ms after the end of saccades is thought to decrease visual sensitivity to saccadic image motion, contributing to visual stability (Ross et al., 2001). Postsaccadic enhancement lasting ~200-400 ms is thought to promote visual perception at fixation (Ibbotson and Cloherty, 2009; Schroeder at el., 2010). Saccadic effects have been reported throughout the visual system, including early cortical and thalamic visual areas (Reppas et al., 2002; Sylvester and Rees, 2006; MacEvoy et al., 2008) as well as in areas of the temporal cortex implicated in visual recognition and memory (Sobotka et al., 1997, 2002; Purpura et al., 2003; Barlett et al., 2011; Monosov et al., 2011; Jutras et al., 2013). Using anatomically-constrained MEG source estimates to measure patterns of cortical activity with high temporal and spatial resolution, previous evidence suggests that visual and central effects of saccades also alter the dynamics of word-evoked cortical activation (amplitude, time course) at multiple stages of visual word processing (Temereanca et al., 2012). It is not known, however, if central saccadic signals impact higher stages of visual word representation in occipitotemporal and/or temporal cortex above and beyond preceding effects in occipital cortex.

Here we use anatomically-constrained MEG to examine cortical activity following the onset of saccades as healthy human subjects engaged in a one-back word recognition task. In a parallel experiment in the same participants, this activity was compared with activity following the onset of external visual stimulation that mimicked saccades. With retinal stimulation similar between these conditions, response differences provide insights into the contributions of central saccadic mechanisms to cortical activity after the onset of saccades, when central saccadic signals are present, vs. after external image motion, when such saccadic signals are absent. We introduced a new approach that combines several procedures to eliminate saccadic ocular artifacts from the MEG signal, an inherent difficulty in active reading research with electrophysiological measurements (Berg and Scherg, 1991; Dimigen et al., 2011; Carl et al., 2012). Cortical regions impacted by central saccadic influences were established using cluster-based analysis across space and time, and results were tested on an independent subset of data using parametric statistics. Employing this approach, we compared cortical activity following the onset of saccades and spanning fixation before word appearance vs. similar external visual stimulation. We then examined whether differential responses colocalized with word repetition priming effects, as well as with known saccadic effects on responses to words (Temereanca et al., 2012). Together, our results provide the first evidence for central saccadic influences in a left-lateralized language network in occipitotemporal and temporal cortex above and beyond saccadic influences at preceding stages of information processing during visual word recognition.

## 2. MATERIALS AND METHODS

### 2.1. Subjects

Participants’ approval was obtained and informed consents were signed before each measurement. Seven healthy right-handed adults (mean age of 29 years, range 23-42 years, 5 males) underwent two MEG sessions for Experiments 1-2, and also a structural MRI scan. All experimental protocols were approved by the Massachusetts General Hospital Institutional Review Board, and procedures were conducted in accordance with the approved guidelines.

### 2.2. Experiment 1 (Natural Saccades)

As described in detail in Temereanca et al, 2012, during a one-back word recognition task, subjects waited for an auditory go-cue at the beginning of each trial to make a saccade between two fixed strings of five crosshairs, 10 degrees apart (Fig. 1A). Saccades were detected in real time using the horizontal electrooculogram (EOG) signal and triggered the subsequent word appearance at the new fixation either 76 ms (early word presentation condition) or 643 ms (late condition) later. These latencies include a fixed delay of 33 ms between the stimulus trigger pulses sent by the Presentation program and the stimulus appearance on the projection screen. These latencies ensured in individual subjects that words appeared at re-fixation only after the end of saccades (see online saccade detection and offline computations below), allowing control of stimulus timing (onset and duration) across conditions. The word stimuli were five-letter novel words (50%) and one repeated word presented for 250 ms. No-word trials were included wherein a string of five Xs presented 1243 ms after the saccade detection marked the end of trial. No-word and late word presentation trials were examined here to evaluate the brain activity evoked by saccades in the absence of words. Early, late and no-word trials appeared in pseudorandomized order, with 1300-1500 ms interstimulus interval. Subjects were instructed to read the stimulus silently and respond as accurately and quickly as possible by pressing a button with their right index finger if the stimulus was the same as that in the previous trial (10%, match trials), and another with their left finger otherwise (90%, non-match trials). We collected 110 trials per condition for each of 10 conditions (early vs. late word presentation, novel vs. repeated words and no-word trials, for right as well as left saccades) in 20 blocks, with short 1-3 min. breaks between blocks and a total recoding time of 90 min. Two additional blocks were used to familiarize the subject with the task before recordings. During recordings, subjects rested their upper jaw on a custom-made bite-bar while comfortably leaning their head against the back of the dewar’s helmet; this approach maintained a steady position of the head relative to the MEG sensors within as well as across recording sessions.

**Figure 1.**
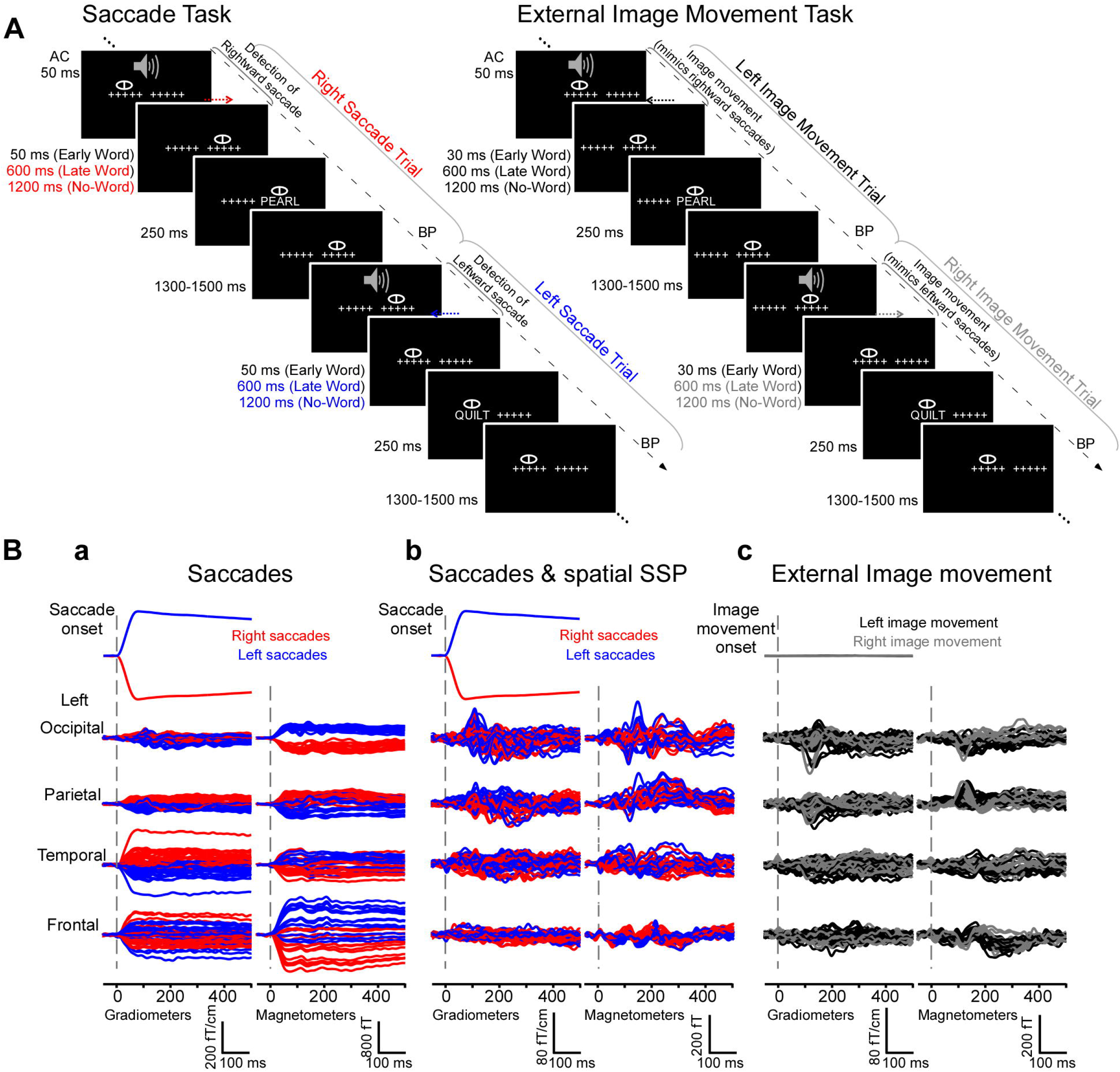
Experimental design and MEG waveforms from an individual subject A, Experiment 1 (natural saccades, left). Subjects performed a one-back word recognition task while reading words presented foveally after saccades detected in real time. Cued by a brief tone, subjects made saccades between two strings of five crosshairs separated by 10° and subsequently maintained fixation before words appeared early or late after saccade detection, or until the end of trial (no-word trials). No-word and late word presentation trials were used to examine cortical activity following the onset of saccades in the absence of words, during a time-window of -200 to 500 ms relative to the onset of saccades. Experiment 2 (external image movement, right). In parallel experiments, the same subjects performed the one-back word recognition task while reading words presented foveally after external image movement that mimicked saccades. The experimental design, including image movement and word presentation timing, matched those in the saccade task. No-word and late word presentation trials were used to examine cortical activity after the onset of saccade-like external image movement. AC, Auditory cue; BP, button press. B, Mean horizontal EOG and MEG waveforms from a representative subject generated by right (red) and left (blue) saccades (a) before and (b) after saccadic artifact reduction with Spatial Signal Space Projection (SSP) method. Right and left saccades produced ocular artifacts with opposite polarity, measured as significant correlations between the horizontal EOG and MEG signals (see Results). SSP filtering eliminated or greatly attenuated artifacts across individual MEG channels. c) Mean horizontal EOG and MEG waveforms generated by left (black) and right (gray) image movement that mimicked right and left saccades, respectively.

Word stimuli were five-letter words balanced across conditions with respect to lexical frequency (Kucera and Francis 1967; range, 1 - 192/million), concreteness index (range, 220 - 648), and stress. Both novel words (50% of trials) and one repeated word were presented. For the repeated word condition, we repeatedly presented a single word either early or late after right and left saccades in order to reliably assess postsaccadic effects on responses in early visual areas that are sensitive to the visual attributes of the stimulus (Temereanca et al., 2012). Analysis of novel vs. repeated word contrast was reported and discussed extensively in Temereanca et al., 2012, revealing left-lateralized word repetition effects consistent with well-established results in language research.

Stimuli were presented on a computer-driven projection and subtended 5 degrees visual angle; there was about 1 letter per degree of visual angle. The whole projection screen subtended 47 degrees.

Occasionally, the electronic circuit did not detect a saccade and as a result failed to trigger the word appearance. For these trials (< 10% of all trials), feedback was provided immediately by the appearance of the word ‘error’ at the missed saccade target location, which cued the subject to correct gaze by fixating the missed location and await a new trial. This small failure rate indicates that subjects were able to perform the saccades consistently and stereotypically.

Cortical responses related to words were reported in Temereanca et al., 2012. In the present study we examined cortical activity following the onset of saccades in the absence of words, estimated from no-word trials and late word presentation trials. We then compared this pattern with the cortical activity following the onset of external visual stimulation that mimicked saccades estimated in Experiment 2.

### 2.3. Experiment 2 (Simulated Saccades)

In parallel experiments words were presented after external image movement that mimicked an eye movement (Fig. 1A). Following the auditory cue, subjects were instructed to maintain their gaze stationary in the center of the screen while the two strings of five-crosshairs 10 degrees apart were moved to mimic the retinal image motion during an eye movement. Words were presented at fixation either early (59 ms) or late (626 ms) after the external image movement offset. No-word trials were also included wherein a string of five Xs presented 1243 ms after movement offset marked the end of trial. These no-word trials as well as the late word presentation trials were examined here to evaluate the brain activity following the onset of saccade-like image motion in the absence of words. Word and no-word trials appeared in pseudo-randomized order, with 1300-1500 ms interstimulus interval. Based on data in Experiment 1, for each subject we computed the mean and variance of the saccade onset latencies with respect to the auditory cue (see offline computations of saccade times below). Random numbers following this distribution were generated and used here to set the onset time of the image movement relative to the auditory cue. Motion velocity and duration matched the average values obtained for saccades in preliminary experiments. Specifically, motion stimuli consisted of a sequence of five frames presented at 60 Hz, which changed the location of the two crosshairs on the screen by a total of 10 deg to mimic the motion stimulus on the retina during 10 deg horizontal saccades (see Figure 1A). The duration of the motion stimulus was 83 ms (5 frames x 1/60 Hz = 83 ms) and matched the average saccade duration of ~80 ms obtained in preliminary experiments; the shift for each individual frame was chosen to match the velocity profile of saccades (as inferred from the EOG signal): 1.5 deg for first frame, 2.6 deg, 2.9 deg, 2.2 deg and 0.8 deg for subsequent frames, respectively. Experiment 2 paralleled Experiment 1 in every other aspect regarding word stimuli, inclusion of no-word trials, number of trials per condition and number of blocks, collection of behavioral data as well as task instructions.

### 2.4. MEG Recordings

Whole-head MEG was recorded in a magnetically and electrically shielded room (Imedco AG, Hägendorf, Switzerland) using a Neuromag Vectorview system (Elekta Neuromag Oy., Helsinki, Finland) with 306-channels arranged in triplets of two orthogonal planar gradiometers and a magnetometer. The acquisition band-pass was 0.01 – 200 Hz and the data were digitized at 600 samples/s. The horizontal and vertical components of eye-movements were recorded concurrently with MEG using two pairs of bipolar EOG electrodes. For subsequent coregistration with the structural MRI and to record the position of the head relative to the sensor array, the locations of four head-position indicator (HPI) coils attached to the scalp, three fiducial landmarks (nasion and auricular points), and additional scalp surface points were digitized using a 3Space Fastrak system (Polhemus, Colchester, VT) integrated with the Vectorview system.

### 2.5. Saccade Detection

Saccades were detected in real time using the EOG signal for horizontal eye movements, which was sent online to a saccade-detection circuit employing filters and a threshold comparator (Temereanca et al., 2012). Saccade detection triggered the word presentation at two delays adjusted in preliminary experiments so that the display changed at refixation only after the end of the saccade, allowing control of word stimulus timing (onset and duration) across conditions.

The times of saccades and word presentation relative to saccades were confirmed offline. The beginning and endpoint of saccades were computed for each subject and experimental condition (e.g., separately for early, late and no-word conditions corresponding to right saccades and left saccades) based on the EOG signal averaged across trials aligned to the time of online saccade detection, using inhouse software in MATLAB (Mathworks, Natick, MA). The algorithm started at peak velocity computed as the first temporal derivative of the EOG trace and searched backward and forward to fixation. Saccade onset and offset were defined as the first point in time preceding peak velocity and the last point following peak velocity, respectively, for which velocity was larger than 3.3 SDs from the mean baseline value (p < 0.001).

These offline computations revealed saccade latencies of 297.1 ± 21.2 ms (mean ± SD) following the auditory cue. They also showed that the saccade-detection circuit detected saccades in the EOG signal 47 ms on average following saccade onset; detection times varied across subjects and for right vs. left saccades, accounting for words appearing on average 21 ms earlier after the end of right than left saccades (range, 14-30 ms) (Temereanca et al., 2012). These offline computations confirmed that words appeared in the early condition 119.9 ± 2.9 ms (mean ± s.e.) and in the late condition 686.7 ± 3.0 ms after the onset of saccades; these times include a fixed delay of 33 ms between the stimulus trigger pulses sent by the Presentation program and the stimulus appearance on the projection screen, which is attributed in part to the projector’s refresh rate of 60 Hz. These computations revealed that right and left saccades had similar durations of 82.8 and 81.3 ms on average, respectively (paired t-test, p > 0.5; range, 75-91 ms).

### 2.6. MEG Data Analysis

#### Correction for Ocular Artifacts, Including Horizontal Saccadic Eye Movement Artifacts

Noisy MEG channels were identified by inspection of raw data and offline averages, and were excluded from subsequent analysis. MEG data were low-pass filtered at 40 Hz. In Experiment 1, trials including right or left saccades detected in real-time were rejected based on large vertical EOG (>150 μV) indicative of blinks. The raw EOG signals aligned to the saccade onsets revealed similar amplitudes and shapes across trials, consistent with subjects performing stereotypical saccades between the two strings of crosshairs. In Experiment 2, trials during steady eye fixation were rejected based on large vertical and horizontal EOG signals (>150 μV) indicative of blinks and horizontal eye movements, respectively. The raw EOG signals from individual trials included in further analysis were visually inspected and no evidence was found for systematic eye movements beyond the horizontal saccades detected in real-time in Experiment 1.

Average waveforms relative to the onset of saccades and saccade-like external image motion were obtained separately from no-word trials and from late word presentation trials, as follows. In each subject, averages were computed separately for right and left saccades aligned to saccade onsets (N ~ 65-100 trials), as well as for right and left saccades together. Similarly, averages were computed separately for right and left external image motion aligned to the image motion onset as well as for right and left movement together.

To map the MEG signal onto the cortex and estimate the spatiotemporal cortical activation following the onset of saccades in the absence of horizontal saccadic artifacts, we introduced an approach that combines two complementary analyses. First we averaged an equal number of right and left saccade trials (after randomly eliminating trials from the more numerous condition). In our experimental design, right and left saccades gave rise to ocular artifacts of opposite sign but similar amplitudes (Fig. 1B) that canceled when averaged, thus effectively annulling horizontal saccadic artifacts. Although visual inspection of the EOG signal revealed no systematic eye movements beyond the horizontal saccades included in the experimental design, it is still conceivable that there are differences in eye movement behavior across right and left saccade conditions, producing asymmetric eye movements which could lead to artifacts not canceled by the averaging procedure. To control for an impact of any remaining artifacts, including those introduced by any asymmetric eye movements, in a second analysis we employed spatial and temporal signal-space projection (SSP) methods separately for right saccades and left saccades, respectively, and only afterwards we averaged across right and left saccade trials. SSP projections computed separately for right and left saccades are expected to eliminate or greatly attenuate artifacts across individual MEG channels before averaging, including any artifacts introduced by asymmetric eye movements. Consequently, similar results in these complementary analyses would provide evidence that saccadic artifacts do not impact results in the present study.

Specifically, to suppress saccadic artifacts, we used the signal-space projection (SSP) method implemented in the MNE software (Hamalainen, 1995; Uusitalo and Ilmoniemi, 1997) as well as a temporal filtering method similar to that described in Taulu and Simola, 2006. For SSP, the spatial subspace containing the ocular artifact was estimated from the MEG data. A covariance matrix was computed from concatenated epochs spanning 5 ms before and 90 ms after right and left saccade onsets. A singular value decomposition (SVD) was performed on the covariance matrix separately for magnetometers and gradiometers. One singular vector in each set of sensors, corresponding to the highest singular value, was used to construct a linear projection operator corresponding to the ocular artifact and applied to the MEG data. The noise subspace corresponding to the ocular artifact was thus eliminated.

In the temporal filtering method, a temporal SSP corresponding to the ocular (non-brain) sources was estimated from the EOG data of each non-overlapping time-window of 16 sec. SVD was performed on the normalized EOG in each time window, resulting in a orthonormal set of eigenvectors used to compute the temporal SSP operator, which was then applied to the MEG data, eliminating temporal components corresponding to the ocular artifact.

Both the spatial and temporal SSP approaches have the potential drawback of attenuating brain signals of interest. For the spatial SSP, this dampening can be rigorously assessed by computing the amount by which the signals from any given cortical region are attenuated by the application of the spatial SSP operator. We found that the average of this dampening was below 0.1, suggesting only a minor effect on the MEG signals of interest. Furthermore, in the source analysis we took this dampening correctly into account by applying the spatial SSP operator to the forward solution and noise covariance matrices. Analogously, the temporal SSP procedure dampens any brain signals which are correlated with the EOG data. Since the space of potential brain signal waveforms is unknown, it is impossible to make a conclusive quantitative assessment of this effect. However, the EOG electrodes are relatively far away from most brain regions, with possibly the most frontal parts of the cortex being an exception, and, therefore, we do not expect that the application of temporal SSP seriously distorts the estimated source waveforms. In addition, it is unlikely that the brain signals themselves have waveforms highly correlated with EOG which relates to the ocular activity. This conclusion is further supported by the fact that the source estimates computed from original data with left and right saccades averaged were similar to those resulting from data which had either spatial or temporal SSP applied to them before the averaging procedure.

Saccadic artifacts were measured by assessing the correlation between the horizontal EOG, which mirrors ocular artifacts, and the MEG signal in both original and filtered data (Fig. 1B). In each subject, EOG and MEG data segments from -10 to 70 ms around saccade onset were concatenated across trials, and a correlation coefficient was computed between concatenated data from the horizontal EOG and each MEG channel. Correlation coefficients were mapped to a normal distribution using a Fisher z-transformation and then tested for significance from 0 using a paired t-test across subjects. Focusing on a period around the saccade, this correlation approach detects the presence of the saccadic artifact with high sensitivity regardless of the artifact amplitude.

### 2.7. Structural MRI

MRI recordings (1.5 T Sonata scanner, Siemens Medical Solutions, Erlangen, Germany) consisted of two structural 3D magnetization prepared rapid gradient echo (MPRAGE) scans (TR/TE/TI = 2.73 s/3.31 ms/1 s, flip angle = 7°, 128 x 1.3 mm-thick sagittal slices at an in-plane resolution of 1 mm^2^) and two multi-echo multi flip angle (5° and 30°) fast low-angle shot (FLASH) scans (TR/TE = 20 ms/(1.8 + 1.82 x *n*) ms, *n* = 0-7). The standard MPRAGEs were used for individual cortical surface reconstructions with FreeSurfer (http://surfer.nmr.mgh.harvard.edu) and for registering MEG data to the individual subject’s anatomy (Dale et al., 1999; Fischl et al., 1999a,b). The FLASH sequences were used to compute the inner skull surface for the boundary element model (BEM). This information was then employed in computing the MEG forward solution. Cortical surfaces were inflated to visualize both gyri and sulci and to morph the hemispheres into a sphere for inter-subject registration based on the sulcal-gyral pattern (Fischl et al., 1999a,b).

### 2.8. Anatomically constrained MEG

MEG average signals were further analyzed to estimate the corresponding patterns of brain activity (current sources) across cortical locations and time. MEG measures the magnetic fields generated by post-synaptic currents in the brain. These current sources were estimated using the linear minimum-norm estimate (MNE) approach (Hamalainen and Ilmoniemi, 1994; Dale and Sereno, 1993) and information of the head anatomy obtained from anatomical MRI data using the MNE software (http://www.nmr.mgh.harvard.edu/martinos/userInfo/data/sofMNE.php; Gramfort et al., 2014). The solution space for the estimated currents was constrained to the gray/white matter boundary reconstructed for each individual from the structural MRI, which was subsampled to 4098 dipole elements per hemisphere with ~5-mm spacing (Dale et al., 1999; Fischl et al., 1999a). A forward solution for the source space was computed using a one-layer BEM model (Hämäläinen and Sarvas, 1989). A noise covariance matrix was calculated from 200-ms baseline periods prior to the auditory cue that preceded saccades (Experiment 1) or external image motion (Experiment 2) in individual trials. The noise covariance matrix and the forward solution were used to create a linear inverse operator (Dale et al., 2000) that was applied to the data at each time point, producing time-courses of activity at each cortical location. Current orientations were approximately constrained to be perpendicular to the cortical surface by setting source variances for the transverse components to be 0.6 times the variance of the normal components (Lin et al., 2006). For group analysis, the inverse solutions were registered to the average cortical surface computed across all subjects using an algorithm matching the cortical folding patterns (Fischl et al., 1999b). The current estimate at each cortical location was divided by the estimated baseline variance, resulting in an F-like statistic (Dale et al., 2000). The square root of the F statistic, which is a signal-to-noise ratio estimate, is analogous to a z-score and allows the visualization of results as dynamic statistical parametric maps (dSPM). The dSPM identifies locations where current strength estimates are most reliable based on their signal-to-noise ratio.

### 2.9. Cluster-Based Statistical Analyses

Differences in estimated cortical activity following the onset of saccades vs. saccade-like external image movement in space and time were established using the MEG signals from no-word trials and cluster-based statistics, a nonparametric permutation-based method that inherently corrects for multiple comparisons (Maris and Oostenveld, 2007). Specifically, a paired t-test across subjects was used to compare saccade vs. external image movement conditions at each time point and for each vertex on the cortex. Spatiotemporal clusters were then computed to include spatially and temporally contiguous sources on the cortex and within the 100-500 ms response window that exhibited significant differences at uncorrected p < 0.05. For cluster calculations on the cortex, we computed the adjacency matrix across vertices using Brainstorm (Tadel et al., 2011). For each cluster, the sum of t-score values was calculated across every spatio-temporal point within the cluster. Permutation testing was performed by randomly permuting conditions within subjects. For each permutation, we repeated previous steps of the analysis and then selected the cluster with the maximum sum of t-score values. We used 1000 permutations to obtain a distribution of maximum sums, which constitutes our null distribution. A p-value was then computed for each original (non-permute) cluster by finding the number of times the values in the null distribution were higher than the original cluster t-score; this p-value is corrected for multiple-comparisons across space and time. We selected original clusters that exhibited corrected p < 0.05.

### 2.10. Regions of interest

For clusters of significant differential responses following the onset of saccades vs. external image movement (corrected p < 0.05), we computed the spatial overlap with anatomical labels generated in FreeSurfer as well as with manually-drawn labels from our previous investigation of word-evoked responses in this subject group (Temereanca et al. 2012). As explained in detail in Temereanca et al., 2012, the latter regions of interest (ROIs) have been selected a priori based on their implication in previous studies of visual word recognition, and because they met the study criteria for significant activation in response to visual words. ROIs were represented on the average brain of all subjects. The same ROIs were used for all subjects by automatic spherical morphing of original labels to individual subjects (Fischl et al., 1999b). Regional time-courses of estimated cortical responses for individual subjects and conditions were computed by averaging the absolute current values within an ROI across voxels at each time point.

Results emerging from the cluster-based statistical analysis were confirmed using independent data from late word presentation trials and parametric statistics. Specifically, for an individual cortical region and time interval corresponding to a spatiotemporal cluster and fixed across subjects, we compared activity following saccades vs. saccade-like external image movement using a two-tailed paired t-test.

The approach employed here to compare cortical activity following the onset of saccades vs. saccade-like external visual stimulation was not feasible for the early word presentation trials because of variation in word-onset times relative to the saccade onset/offset for right vs. left saccades and across subjects (see Saccade Detection section above), which generate differences in visual stimulation across conditions.

## 3. RESULTS

### 3.1. Estimated cortical activity related to saccades

#### 3.1.1. Control for artifactual effects of horizontal saccades on MEG signals

Cortical activity following the onset of saccades was examined in seven healthy volunteers engaged in a one-back word recognition task, using anatomically-constrained MEG (see Methods; Fig. 1A). Subjects made horizontal saccades between two strings of five crosshairs 10° apart and subsequently maintained fixation before words (novel or repeats) appeared early (76 ms) or late (643 ms) after saccade detection in real time, or until the end of trial (no-word trials). No-word and late word presentation trials were examined here to evaluate the cortical activity evoked by saccades in the absence of words, during a time-window of -200 to 500 ms relative to the onset of saccades. This activity reflects a combination of the retinal activity during the saccade, the retinal activity at the onset of fixation period, as well as central saccadic signals gated by oculomotor brain regions.

The average MEG responses during -200 to 500 ms were similar for no-word and late word presentation trials, respectively, as expected because of identical visual stimulation and task conditions. Figure 1B illustrates averages of the horizontal electrooculogram (EOG) and MEG responses for right and left no-word trials (N ≈ 90) in an individual subject. Right and left saccades produced ocular artifacts with opposite polarity and similar magnitudes, measured as significant correlations between the horizontal EOG which mirrors ocular artifacts and MEG signals (Fig. 1B). Artifacts were larger over frontal and most anterior temporal sensors and gradually decreased in magnitude over temporal, parietal and occipital sensors.

To remove saccadic artifacts from the MEG data, we averaged a balanced number of right and left saccades, cancelling associated artifacts of similar magnitude but opposite polarity. To control for an impact of potential remaining artifacts attributed to any asymmetric eye movements across right and left trials, in a complementary analysis ocular artifacts were removed from MEG data before the averaging procedure using spatial and temporal Signal Space Projection (SSP) filtering separately for right and left saccade trials (see Methods and Materials). Each of these filtering methods eliminated or greatly attenuated artifacts across individual MEG channels (Figure 1B, b). Remaining artifacts were evaluated by computing the correlation between the horizontal EOG and MEG waveforms around the time of saccades (see Methods). Using this highly sensitive detection method that ignores the amplitude of the remaining artifact, significant correlations were found in 214 out of 306 MEG channels before filtering, but only in 95 frontal channels after spatial SSP. Most importantly, the application of the SSP revealed strong MEG responses with a stable baseline and onset latencies not correlating with the EOG saccade waveform. Averages of right and left saccades were computed for the original signal as well as after artifact removal with spatial and temporal SSP, and produced similar results for all analyses described below, providing evidence that saccadic artifacts do not impact the results reported here.

Estimates of cortical activity were calculated across locations and time using a distributed source modeling approach that constrained current sources to the cortical surface of each participant reconstructed from structural MRI (Dale et al., 1993). Noise-normalized dSPMs (Dale et al., 2000) were computed to evaluate the statistical significance of estimated responses relative to baseline activity measured prior to the auditory cue in each trial. Fig 2 illustrates snapshots of average dSPMs across subjects for two time windows after saccade onset, computed separately for right saccades and for the average of right and left saccades. For right saccades, activity at 60 ms was dominated by saccadic ocular artifacts which exhibited a maximum in the orbitofrontal cortex and extended throughout the frontal and anterior temporal cortex and to a lesser degree to more posterior temporal, parietal and occipital cortex. Spatial and temporal SSP filters greatly reduced ocular artifacts in frontal cortex and temporal poles, and eliminated artifacts in more posterior temporal, parietal and occipital regions. Further, estimates of the average of right and left saccades reliably showed the largest signal-to-noise ratio and minimum ocular artifacts in the most affected regions (orbitofrontal cortex and temporal pole) as well as no contamination across the remaining of the cortical surface. These estimates based on averages of right and left saccades were further analyzed here to examine the cortical activity following the onset of saccades.

**Figure 2.**
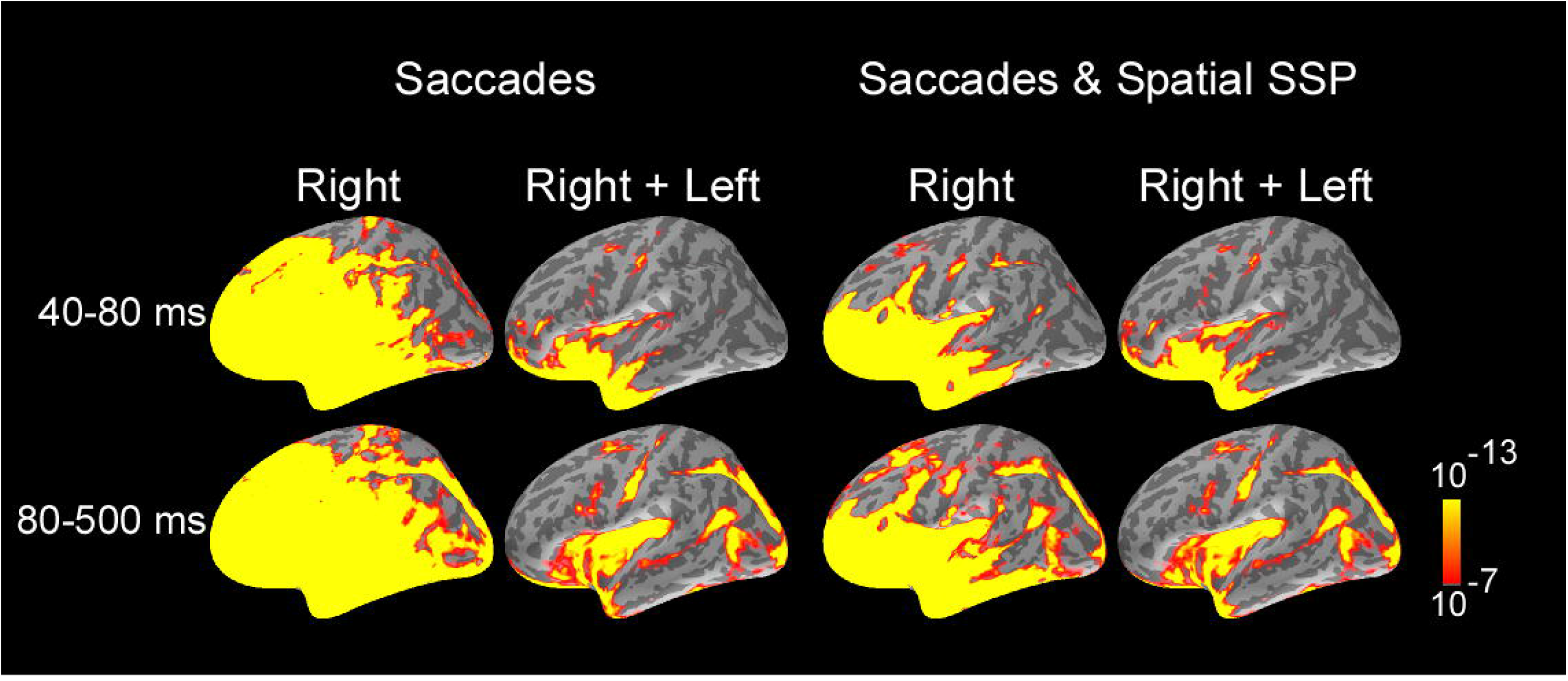
Average dSPMs across subjects and selected response windows following saccade onset, computed separately for right saccades and for the average of right and left saccades. For right saccades, activity at 60 ms was dominated by saccadic ocular artifacts. Ocular artifacts were greatly reduced for MEG data filtered with SSP. Estimates of the average of right and left saccades showed the largest signal-to-noise ratio and no contamination with ocular artifacts across most of cortical surface, and were further analyzed to examine cortical activity following the onset of saccades. Average dSPMs are displayed on the inflated hemispheres of the average brain of all subjects (N=7). Significance is indicated with color bars.

#### 3.1.2. Overall Activity and Time Courses

Figure 3 illustrates the progression of estimated cortical activity following the saccade onset. Activity was prominent in occipital pole and calcarine sulcus starting at ~70 ms, and spread in parallel through the dorsal and ventral visual streams to parietal, temporal and frontal cortices. Specifically, activity engaged posterior temporal cortex including putative motion-sensitive area MT+, and spread to posterior Sylvian fissure, planum temporale and superior insula; and also recruited regions of frontoparietal networks including areas of superior parietal cortex and intraparietal sulcus, as well as the frontal eye field and ventral precentral sulcus. Activity also spread to medial regions including parietooccipital cortex, precuneus and posterior cingulate; and recruited medial temporal and retrosplenial cortex. Within the ventral stream, activity recruited occipitotemporal and inferotemporal regions. Activation in cortical regions exhibited early peaks (range, 80-130 ms) that were followed by subsequent response components (Figs. 3–4) likely reflecting ongoing local processing and long-range network interactions.

**Figure 3.**
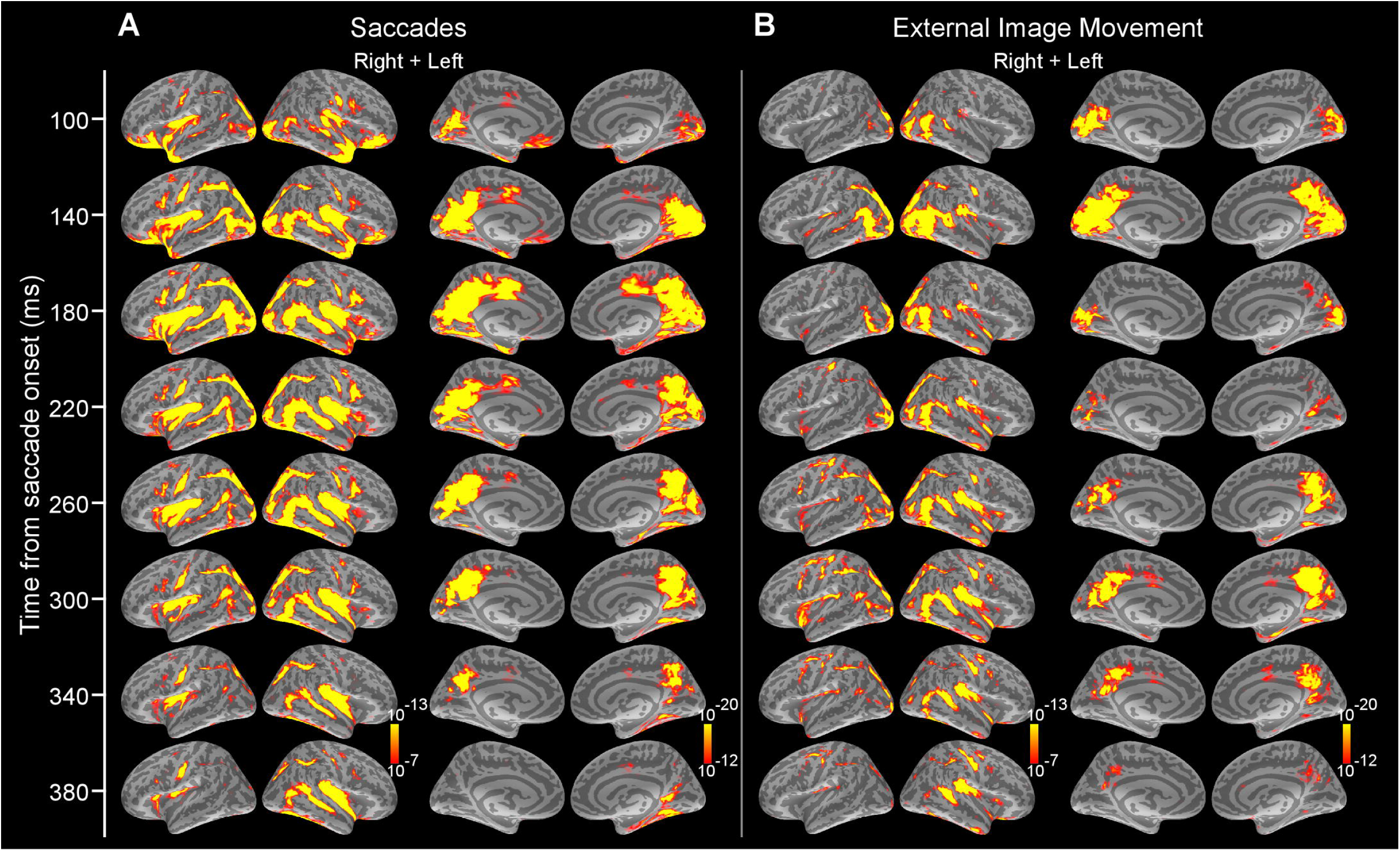
Average dSPMs following the onset of saccades in Experiment 1 and external image movement that mimicked saccades in Experiment 2. A. Experiment 1. Snapshots of average dSPMs at selected latencies after saccade onset. B. Experiment 2. Snapshots of average dSPMs at selected latencies after saccade-like external image motion. Saccades and saccade-like external visual stimulation produced early-latency responses beginning ~70 ms after onset in occipital cortex and spreading through the ventral and dorsal visual streams to temporal, parietal and frontal cortices. Average dSPMs are displayed on the inflated hemispheres of the average brain of all subjects (N=7). Significance is indicated with color bars.

**Figure 4.**
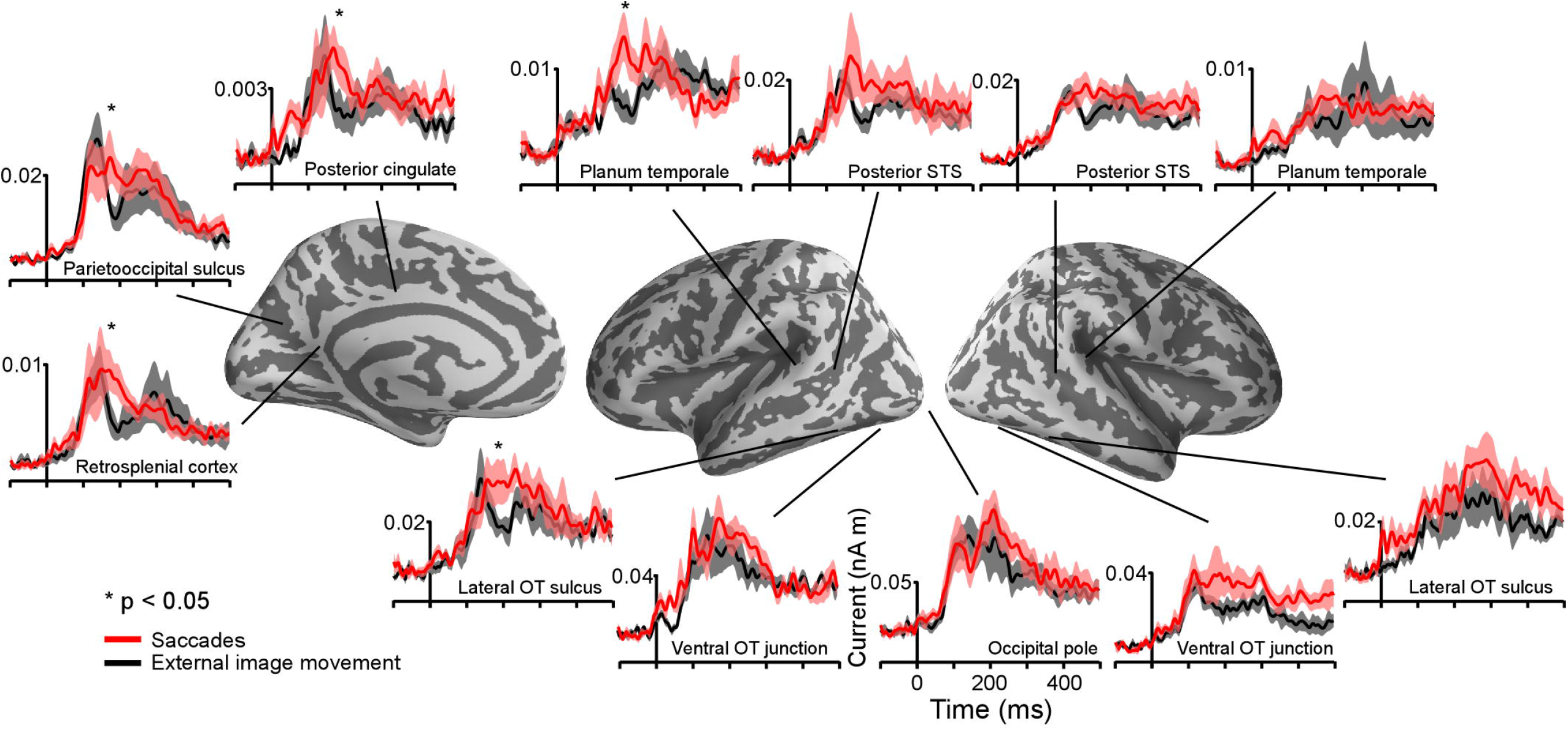
Average time courses of estimated currents in selected cortical regions following the onset of saccades and external image movement that mimicked saccades. Averages were computed across all subjects. Lines are mean responses; shaded areas are mean + SEM. * mark regions and time-courses corresponding to spatiotemporal clusters that exhibited significant differential responses after the onset of saccades versus saccade-like background movement. These clusters are described in Figure 5 and Table 1.

The time course of estimated cortical activity produced by saccades was visualized in regions of interest (ROIs) (see Methods). Regional waveforms, illustrated in Figure 4, were computed in each ROI and individual subject by averaging values across all voxels. Waveforms showed stable baselines and short-latency responses (range, ~70-110 ms) with multiple peaks and a time-course of ~400 ms, which were not correlated with the EOG saccade waveform. Further, regional waveforms estimated based on right and left saccade averages computed from the original data vs. after filtering with spatial and temporal SSP were similar (data not shown), indicating that activation estimates reflect neural sources rather than ocular artifacts (See Methods).

### 3.2. Estimated cortical activity related to saccade-like external visual stimulation

#### 3.2.1. Overall Activity and Time Courses

Similar to the activity produced following the onset of saccades, cortical activity after the onset of saccade-like external visual stimulation produced early-latency responses in occipital pole and calcarine sulcus at ~70 ms and spread to temporal, parietal and frontal regions (Figure 3). Figure 4 illustrates regional waveforms in selected cortical areas. The early phase of the response in occipital pole was remarkably similar for saccades and saccade-like external visual stimulation, reflecting similar retinal stimulation across conditions. Differences in cortical activity following the onset of saccades versus similar external visual stimulation were established using cluster based analysis and then confirmed using parametric statistics as described below.

### 3.3. Comparison of estimated cortical activity related to saccades and saccade-like external visual stimulation

Differences in estimated cortical activity following the onset of saccades and saccade-like external image movement were statistically evaluated using the MEG signal from no-word trials and cluster-based statistical analysis across the whole cortex and response time (see Methods). Specifically, paired t-tests were performed at each time point within 80-500 ms response-window and for each vertex on the cortex. Significant clusters (corrected p < 0.05) with time-courses between 150-350 ms after onset of image motion emerged in several cortical regions illustrated in Figure 5 and summarized in Table 1. Computation of the spatial overlap with anatomical regions segmented in FreeSurfer revealed that significant clusters overlapped with i) left lateral occipitotemporal and nearby inferotemporal cortex, ii) left posterior Sylvian fissure including planum temporale, and iii) medial parietooccipital, posterior cingulate and retrosplenial cortices. Regional responses in these ROIs are illustrated in Figure 4 to visualize the time courses of differential activity across conditions. Activity baselines as well as latencies and earliest phase of responses were similar following saccades and saccade-like external image motion, providing further evidence that current source estimates reflect neural sources rather than saccadic artifacts.

**Figure 5.**
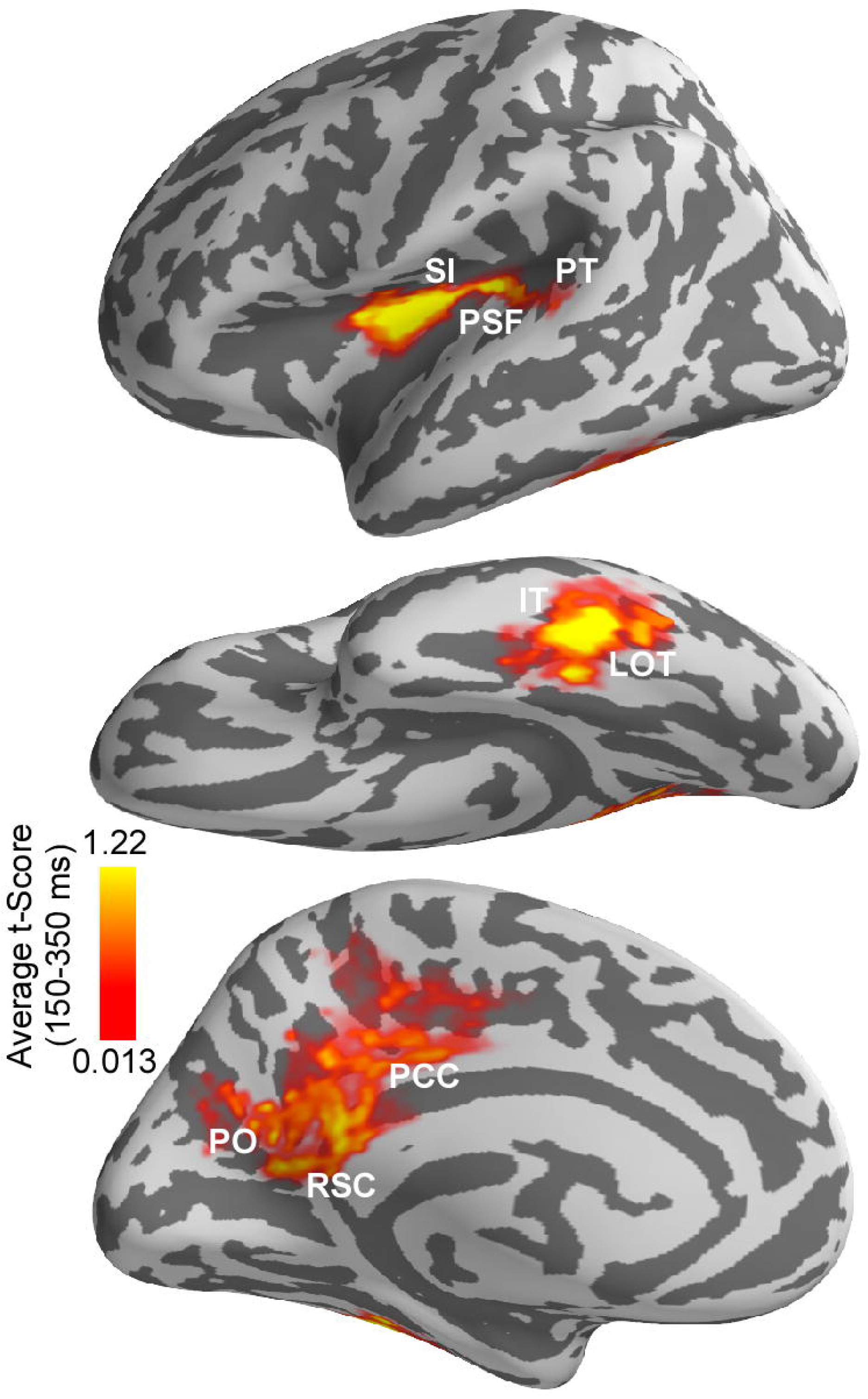
Differences in estimated cortical activity following the onset of saccades vs. saccade-like external image movement. Differences were evaluated using cluster based analysis across whole cortex and response time. Significant spatiotemporal clusters (corrected p < 0.05) with time-courses between 150–350 ms emerged in a left-lateralized cortical network. This network overlapped with anatomically defined (i) left lateral occipitotemporal (LOT) and nearby inferotemporal (IT) cortex, (ii) left posterior Sylvian fissure (PSF), superior insula (SI) and nearby planum temporale (PT), and (iii) medial parietooccipital (PO), posterior cingulate (PCC) and retrosplenial cortex (RSC). Average t-score values over 150–350 ms response-window are illustrated for significant clusters only. Average t-score values are indicated with color bars.

**Table 1.** Results from spatiotemporal cluster-based analysis indentifying cortical regions and tim courses corresponding to significant differences in activity following the onset of saccades versu external image movement that mimicked saccades. Cluster-based analysis was performed across th whole cortex and response time. * corrected p < 0.05.

| Clusters | Time course (ms) |
| --- | --- |
| Lateral Occipitotemporal Sulcus<br>& nearby Inferior Temporal | **156-225\*** |
| Posterior Sylvian Fissure &<br>nearby Superior Insula & Planum<br>Temporale | **171-225\*** |
| Retrosplenial, Posterior Cingulate<br>& Parietooccipital Sulcus | **164-214\*** |

\* $p < 0.05$

Computation of the spatial overlap between the left-lateralized occipitotemporal and temporal clusters and the word repetition priming effects (novel words vs. repeats) reported previously in this population (Temereanca et al., 2012) revealed that they are colocalized. In addition, this left-lateralized network colocalized with saccadic effects on responses to words at fixation, assessed using the early vs. late word presentation contrast (Temereanca et al., 2012).

For the cortical regions corresponding to significant clusters, the presence of differential activity during 155-225 ms was also tested using independent data from late word presentation trials and two-tailed t-tests, and was found significant (paired t-test, all three p’s < 0.03). Further, while significant differential activity within this time window was found in left lateral occipitotemporal cortex (p < 0.03), differences within this time window were not significant in lower visual regions such as occipital pole and left occipitotemporal junction (p’s > 0.05) which provide the incoming afferent information. Also, no differential activity was found in the right lateral occipitotemporal cortex (p > 0.05).

A repeated-measure one-way ANOVA with within-subjects factor of region of interest was conducted to compare the degrees of response change from 155-225 ms following the onset of saccades vs. saccade-like external visual stimulation in three selected ROIs, including occipital pole, left ventral occipitotemporal junction, and left lateral occipitotemporal and nearby inferotemporal cortical cluster. The degrees of response change were significantly different between ROIs (29.7% vs. 42.6% vs.70.0%, F(2,6) = 5.7154, p = 0.018). Pairwise tests revealed that compared with occipital pole, the degree of response change was significantly larger in left lateral occipitotemporal and nearby inferotemporal cortex (29.7% vs.70.0%, p = 0.006), but similar, albeit slightly increased, in left ventral occipitotemporal junction (29.7% vs. 42.6%, p = 0.32), a neighboring visual region processing afferent visual signals from occipital cortex.

## 4. DISCUSSION

Using a new approach to eliminate saccadic ocular artifacts from the MEG signal, the present study examined the spatiotemporal pattern of cortical activity following the onset of saccades as participants engaged in a one-back word recognition task. To evaluate extraretinal, central saccadic influences, the activity produced after the onset of saccades and spanning postsaccadic fixation before word appearance was compared with cortical activity following the onset of external visual stimulation that mimicked saccades. Results revealed robust differential activity in a left-lateralized cortical network during 150-350 ms response-window spanning fixation. Specifically, in line with previous research, differential activity overlapped with anatomically-defined medial parietooccipital, retrosplenial and posterior cingulate cortex, brain regions known to be implicated in visuospatial orientation, gaze self-monitoring and working memory (Haarmeier et al., 1997; Tikhonov et al., 2004; Binder et al., 2009). In addition, differential activity overlapped with left lateral occipitotemporal and nearby inferotemporal cortex implicated previously in visual recognition, including visual word-form access (Tarkiainen et al., 1999; Solomyak and Marantz, 2009; Dehaene and Cohen, 2011); and left posterior Sylvian fissure and nearby multimodal cortex implicated previously in self-induced visual motion perception and language function (Thier et al., 2001; Wise et al., 2001). This left-lateralized network colocalized with word repetition priming effects as well as with known saccadic effects on responses to words presented at fixation (Temereanca et al., 2012), suggesting that central saccadic signals influence visual word processing in these regions. As discussed below, the present results could account for the pattern of various degrees of modulation across cortical regions produced by saccades on subsequent responses to words reported in Temereanca et al., 2012. Together, results provide the first evidence for central saccadic influences in a left-lateralized language network in occipitotemporal and temporal cortex above and beyond saccadic influences at preceding stages of information processing during visual word recognition.

### 4.1. Dynamic patterns of cortical activity related to saccades

Examining MEG waveforms around saccades is challenging due to ocular artifacts that obscure brain signals (Fig. 1; Carl et al., 2012). In our previous report of cortical responses to words presented after saccades, we eliminated saccade-related ocular artifacts by subtracting the MEG signals generated by saccades alone, which include the eye-movement artifact and brain activity associated with saccades (Temereanca et al., 2012). Here we used a new approach to estimate the cortical activity following the onset of saccades in the absence of words at fixation, employing two complementary analyses. Firstly, we averaged an equal number of right and left saccades, which effectively canceled artifacts with similar waveforms but of opposite polarity. In a second approach, using spatial and temporal SSP methods, we eliminated or significantly dampened ocular artifacts across MEG channels and only then averaged across right and left saccades. We conducted our analyses both for averages of right and left trials computed from the original data in the absence of SSP and with the SSP methods followed by this averaging procedure, and obtained similar results, which provides evidence that estimates reflect neural activity rather than contamination from ocular artifacts. Using these complementary analyses, we present here a first report of the spatiotemporal pattern of MEG estimated cortical activity following the onset of saccades.

Consistent with previous electrophysiological recordings of eye movement potentials in monkey (Purpura et al., 2003), saccades evoked a short-latency visual response ~70 ms after onset in occipital cortex. In addition, we found that the earliest occipital response components were virtually the same for saccades and saccade-like external visual stimulation, suggesting similar early processing of closely matching self-induced (saccadic) vs. external image motion. The visual response evoked by saccadic image motion, albeit not consciously perceived, may impact information processing at fixation via interactions between visual signals during and after saccades (Ibbotson and Cloherty, 2009; Temereanca et al., 2012). Later components of occipital responses exhibited differences across conditions that were not statistically significant using our analysis, although likely they reflect central saccadic signals known to modulate occipital activity during and after saccades (e.g., Sylvester and Rees, 2006; Temereanca et al., 2012).

Cortical activity following the onset of saccades spread in parallel through the ventral and dorsal visual pathways and encompassed temporal, parietal, and frontal cortices, in agreement with previous human functional neuroimaging and monkey electrophysiological data. For example, consistent with studies of sensory-motor pathways, activity propagated from early visual cortex to motion-selective area MT+ (Tootell et al., 1995) as well as to superior parietal lobule, lateral intraparietal sulcus (LIP), frontal eye field (FEF), and ventral precentral sulcus, which are component regions of frontoparietal networks implicated in visual target selection, attention and saccade planning and execution (Colby and Goldberg, 1999; Yeo et al., 2011). Recent fMRI studies reported activation in these frontoparietal regions during natural text reading, consistent with a similar role in controlling eye movements in reading (Choi et al, 2014; Choi and Henderson, 2015). In addition, activity encompassed distributed cortical regions associated with visual, memory, and language functions, including midline precuneus and posterior cingulate cortex subserving visuospatial orientation and gaze self-monitoring; retrosplenial cortex and medial temporal regions supporting working memory (e.g., Mesulam, 1990; Yeo et al., 2011); and occipitotemporal and inferotemporal cortices implicated in visual recognition, including visual word recognition in reading. Overall, our new approach that employs anatomically-constrained MEG and cluster-based statistics opens up possibilities for future studies of the temporal relationship and putative interactions between visual, language, working memory and motor processes during complex behaviors such as natural reading.

### 4.2. Cortical regions and time-courses with differential activity following saccades vs. similar external visual stimulation

Comparison of cortical activity following the onset of saccades vs. saccade-like external visual stimulation during one-back word recognition uncovered differential responses with time-courses between 155-225 ms in several cortical regions previously associated with distinct functions. Specifically, robust differential activity was found in parietooccipital cortex and posterior Sylvian fissure, consistent with previous monkey electrophysiology and human lesion studies that implicate these regions in the perception of self-induced image motion (Haarmeier et al., 1997; Thier et al., 2001; Tikhonov et al., 2004). Interestingly, a network of interconnected regions at and near posterior Sylvian fissure that includes superior insula, ventral somatomotor cortex, and auditory cortex has been implicated in other functions critical for self-monitoring and language such as processing speech movements and hearing one’s own voice (Wise et al., 2001; Yeo et al, 2011).

In addition, we found differential responses in left lateral occipitotemporal and nearby inferotemporal cortex. Since differences in occipital regions were significantly smaller, the emergence of disproportionately large, robust differences in left lateral occipitotemporal cortex, the recipient of afferent visual signals from occipital regions, suggests central saccadic influences in this cortical area above and beyond saccadic influences in occipital cortex. Alternatively, it is possible that small response differences in occipital cortex are amplified by the local circuitry in left lateral occipitotemporal cortex, without additional input from a saccade-related mechanism. However, this possibility alone is less likely since evidence from our previous study (Temereanca et al., 2012), discussed below, indicates different degrees of response changes in occipital and lateral occipitotemporal cortex following saccades compared to saccade-like external visual stimulation.

Convergent evidence in our study supports a role of this latter differential activity in visual word processing. Firstly, we found that this differential activity is left-lateralized and colocalized with left-lateralized word repetition priming effects that are typically employed to study visual word processing. In addition, this differential activity also colocalized with saccadic modulation of word-evoked responses reported in Temereanca et al., 2012. The left lateral occipitotemporal and nearby inferotemporal cortices have been implicated previously in word-form access and lexico-semantic processing, respectively (Halgren et al., 1994; Dehaene et al., 2002; Vinckier et al., 2007; Chan et al., 2011). We also found differential responses with a similar time-course overlapping left planum temporale in Sylvian superior temporal cortex, an area previously implicated in grapheme-to-phoneme coding and known to be functionally coupled to the ventral word-form system (Dehaene et al., 2010; van der Mark et al., 2011).

The present results may explain the varying sizes of effects across cortical regions produced by saccades and external image-motion on subsequent word-evoked responses reported in Temereanca et al., 2012. This previous paper, summarized in Figure 6A, revealed reduced behavioral performance and responses to words presented early versus late after saccades as well as early versus late after external image movement in occipital pole, left ventral occipitotemporal junction and lateral occipitotemporal cortex, suggesting that visual effects attributed to image movement during saccades suppress subsequent word responses at re-fixation. Importantly, comparison of saccadic and external movement effects revealed more pronounced word-response modulations after saccades in occipital pole and left ventral occipitotemporal junction, suggesting influences of central saccadic signals in addition to visual influences of image movement. Further, these effects changed in left lateral occipitotemporal cortex, reflecting a boost in the word response immediately after saccades but not after external image-motion. The present results suggest that this change reflects further modulation of word-evoked responses in left lateral occipitotemporal cortex attributed to central saccadic influences (Fig. 6B). Together with previous results, our new findings suggest that central saccadic signals modulate visual word processing in left lateral occipitotemporal and nearby inferotemporal cortex above and beyond saccadic influences at preceding stages of information processing in lower visual areas.

**Figure 6.**
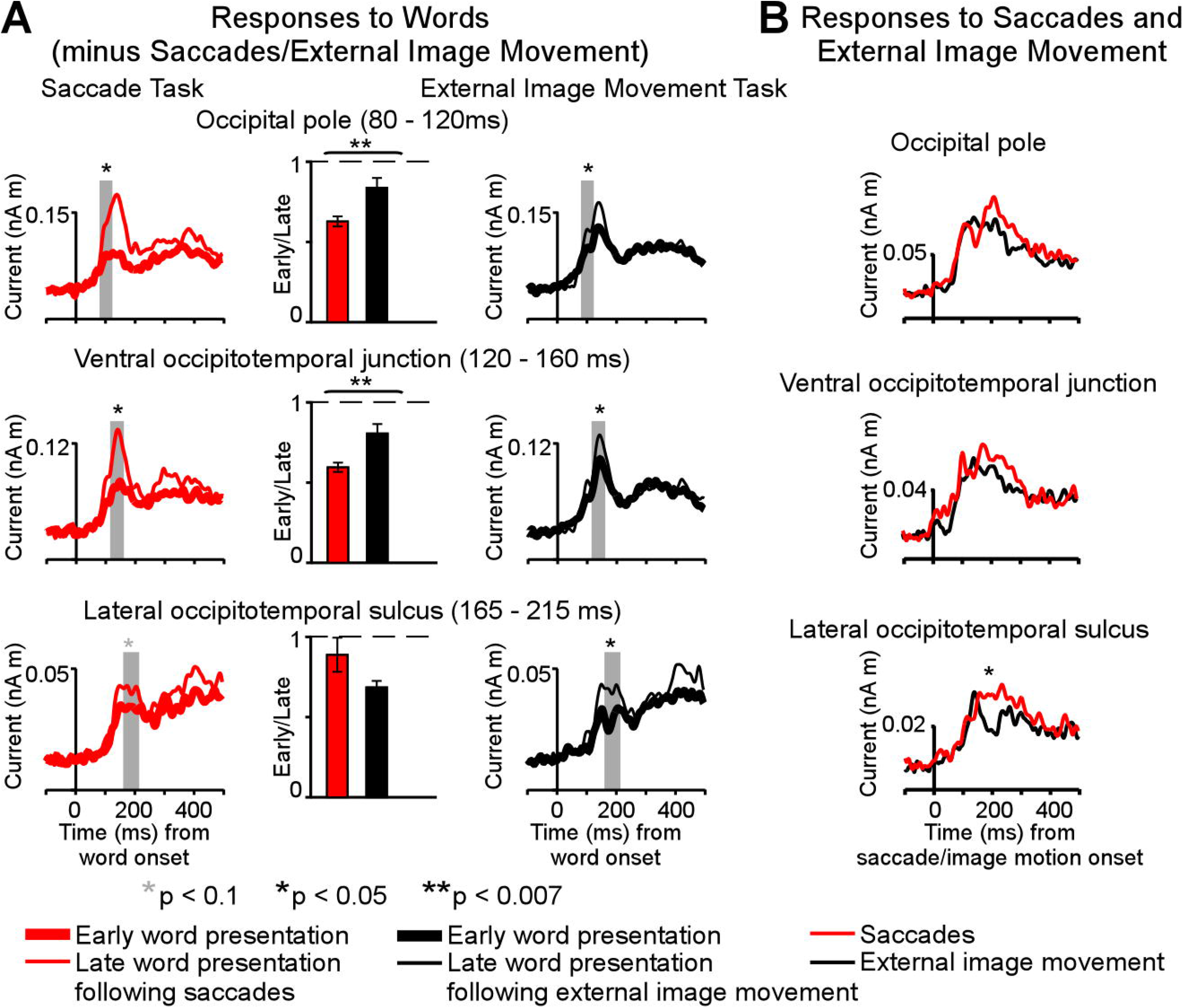
Summary of saccadic contributions to cortical activity during a one-back visual word recognition task. A. Differential effects of saccades and external image movement on word-evoked responses (modified from Figure 6, Temereanca et al., 2012) Cortical responses were reduced to words presented early versus late after saccades, as well as to words presented early versus late after external image movement that mimicked saccades, suggesting that retinal motion contributes to postsaccadic inhibition. Further, response attenuation was significantly larger in the saccade than external image movement condition in occipital pole (80-120 ms) and ventral occipital junction (120-160 ms), consistent with central postsaccadic influences on word processing in these regions. In contrast, the degrees of response modulations in left occipitotemporal cortex (165-215 ms) were similar across conditions, suggesting that central postsaccadic effects very across cortical regions. In particular, these effects on word-evoked responses changed in left lateral occipitotemporal cortex, reflecting a boost in the word response immediately after saccades but not after external image-motion. B. Cortical activity following the onset of saccades and saccade-like external image movement in the absence of words, in the same cortical regions as above. Robust differential activity following th onset of saccades versus similar external visual stimulation emerged during 150-350 ms in left later, occipitotemporal cortex. Since differences in lower occipital regions were smaller, below statistic! significance in our cluster analysis, the emergence of disproportionately large differences in left later occipitotemporal cortex, the recipient of afferent visual signals from lower occipital regions, sugges central saccadic influences in this cortical area above and beyond saccadic influences in occipital corte Together with previous results, our new findings suggest that central saccadic signals modulate visu word processing in left lateral occipitotemporal cortex above and beyond saccadic influences preceding stages of information processing in occipital cortex.

### 4.3. Saccadic contributions to reading

Natural reading typically involves sequences of frequent, small (1-2°) saccades on target words embedded in text, and therefore differs from our experimental conditions in a number of ways, including the size of saccades, the presence of parafoveal information, and perceptual and cognitive processing difficulty. Human EEG measurements during reading (Kornrumpf et al., 2016) and monkey electrophysiology in occipital cortex during active vision (MacEvoy et al., 2008) suggest that saccadic signals interact with these factors and therefore may impact information processing in reading to a different degree and with a different time course than in our experimental paradigm. The present results isolate effects of the saccade itself in occipitotemporal extending to nearby inferior temporal cortex, where they predict interactions between saccadic signals, attention and reading processes, including these factors.

Our findings complement active vision research that has revealed central saccadic signals at various levels of visual representation in the ventral stream. Electrophysiological recordings in primates have revealed central saccadic effects in thalamus and a number of visual cortical areas, including a biphasic modulation of visual responses consisting of transsaccadic suppression followed by postsaccadic enhancement (Reppas et al., 2002; Ibbotson and Krekelberg, 2011) as well as changes in oscillatory activity (Purpura et al., 2003; Rajkai et al., 2008). Central saccadic signals also modulate single-unit and local field potential (LFP) visual responses, functional connectivity, and oscillatory phase in regions of the temporal lobe in monkey, suggesting a role in high level visual recognition, memory processes and their coupling (Sobotka et al. 1997; Sobotka et al., 2002; Barlett et al., 2011; Jutras et al., 2013). The timing and frequency properties of saccade-triggered LFP signals recorded simultaneously in occipital and temporal regions suggest that the nature of saccadic modulation varies across the cortex (Purpura et al., 2003). The present results are consistent with these previous findings, supporting the presence of distinct saccadic contributions to visual word processing in occipital and temporal cortex, respectively. These saccadic contributions are likely mediated by distributed occulomotor regions that control eye movements in both scene viewing and text reading (Choi et al, 2014; Choi and Henderson, 2015).

Because the retinal stimuli analyzed here consisted of two strings of five-crosses lacking orthographic-lexical attributes, our results suggest a task-dependent (rather than stimulus-driven) mechanism impacting the left occipitotemporal cortex during visual word recognition. In the present experiment, the task goals were identical across conditions and likely involved similar task strategies. It is possible, however, that central saccadic signals interact with attentional mechanisms, impacting processing at refixation. Eye movements and attention are controlled by overlapping neural circuits and have been proposed to interact during active vision (Ross et al., 2001; Purpura et al., 2003). Selective attention is known to enhance processing at various levels of visual word representation depending on task goals (Ruz and Nobre, 2008), and could contribute to activity in left occipitotemporal cortex during our one-back word recognition task. An interaction between central saccadic signals and attention could produce differential modulation of information processing in left occipitotemporal cortex following saccades, when central saccadic signals are present, vs. after similar external image motion, when such saccadic signals are absent.

Another possibility explaining our results is that saccadic contributions to visual word processing vary with reading experience known to alter cortical organization and function. Indeed, literacy is known to enhance left occipitotemporal and planum temporale activations evoked by written and spoken language, and their coupling (Carreiras et al., 2009; Dehaene et al., 2010). Further, literacy appears to mobilize dorsal stream mechanisms for visual motion perception as evidenced by enhanced visual motion function (V5/MT activity and coherent motion sensitivity) in good vs. poor readers and in typical readers vs. dyslexic individuals, whereas visual motion is similar for individuals matched on reading ability (Boets et al, 2011; Olulade et al., 2013). Thus, literacy may shape brain activations evoked by saccadic and external visual motion, and the strength of central saccadic influences in left occipitotemporal cortex and planum temporale, regions undergoing critical changes during reading acquisition. Understanding saccadic contributions during reading and how they change with reading experience may provide important insights into the complex interplay between eye movements and visual, language, attention and memory processes underlying reading and its disorders.

## AUTHOR CONTRIBUTIONS

YC, SK, and STI conceived and designed the current study. YC, SK, ST, and STI performed the analyses for the study. YC, SK, ST, GK, EB, MH, and STI wrote, edited and revised the manuscript.

## ACKNOWLEDGMENTS

We thank Drs. Seppo Ahlfors and Annika Hulten for thoughtful comments on the manuscript. This work was supported by NIH grants R03HD050627 (STI), DP1OD003646 (EB), R01EB006385 (EB and MH), R01EB009048 (MH), National Institute of Biomedical Imaging and Bioengineering 5P41EB015896 (MH), and a Claflin Distinguished Scholars Award (STI) and Executive Committee on Research (ECOR) Award (STI) from Massachusetts General Hospital. The authors declare no competing financial interests.

